# bMSI-CAST: a systematic method for next generation sequencing-based microsatellite instability detection in plasma cell-free DNA

**DOI:** 10.1101/2021.02.22.432191

**Authors:** Fengchang Huang, Lili Zhao, Hongyu Xie, Jian Huang, Xiaoqing Wang, Jun Yang, Yuanyuan Hong, Jingchao Shu, Jianing Yu, Qingyun Li, Hongbin Zhang, Weizhi Chen, Ji He, Wenliang Li

## Abstract

Microsatellite instability (MSI) is a well-established prognostic and predictive biomarker in certain types of cancers. MSI detection using tumour tissue is often limited by the availability of specimens. Next generation sequencing (NGS)-based MSI detection in plasma cell-free DNA (cfDNA) is challenged by a much lower signal-to-noise ratio. We developed a highly accurate cfDNA MSI detection method called bMSI-CAST (blood MSI Caller Adjusted with Sequence duplicaTes), with improvement on three features including a set of locus selection principles ensuring loci with high robustness and compatibility across sequencing platforms, an MSI-specific duplicate removal strategy, and a calling algorithm that dynamically matches baselines with a broad range of duplication levels. Analytical validation via MSI-high (MSI-H) cell gDNA showed an LOD of 0.15%. Furthermore, in an analysis of 95 evaluable cfDNA samples from patients with gastrointestinal cancers, bMSI-CAST exhibited a positive predictive agreement (PPA) of 92.9% (39/42) and negative predictive agreement (NPA) of 100% (53/53) with tissue MSI-PCR. In conclusion, bMSI-CAST provides novel and advanced solutions to key aspects fundamental to cfDNA MSI calling but not sufficiently addressed by existing methods, and it is a validated method ready to be applied to aid clinical decisions for cancer patients.

## INTRODUCTION

Microsatellite instability (MSI), a phenotypic manifestation of DNA mismatch repair (MMR) deficiency, is an emerging National Comprehensive Cancer Network-recommended biomarker for the diagnosis and the prognostic evaluation of patients with colorectal cancer (1,2) and other solid tumours (3). In clinical settings, assessment of MSI status in tumour tissue is limited by the availability of high-quality specimens; non-invasive cell-free DNA (cfDNA) assays have been developed to circumvent this issue and help expand the clinical application of MSI analysis. Recently, several cfDNA-based MSI detection methods have been clinically applied to predict patient response to immune checkpoint inhibitors (4,5). Compared to electrophoresis-based MSI-PCR, which evaluates only a few microsatellite loci, or immunohistochemistry (IHC), which indirectly assesses a smaller number of MMR proteins (6), next-generation sequencing (NGS)-based MSI assays can assess hundreds or thousands of microsatellite loci, lending the necessary statistical power for accurate MSI detection in samples with low cancer cell fractions.

NGS-based MSI calling algorithms that contrast the repeat lengths of alleles or between a test sample and normal baseline (7-10), employed by tissue-based MSI methods, have been extended to cfDNA MSI detection, but analysis of cfDNA poses a greater challenge in extracting signals from background noise generated during PCRs as the cancer content fraction is typically three or four magnitudes lower (5). Technically, this requires not only improvements in experimental techniques, such as increasing sequencing depth and using unique molecular identifiers (UMIs), but also more sophisticated algorithms to discriminate true instability from noise or errors introduced during PCRs.

To date, a group of recently developed plasma cfDNA-based MSI detection methods (5,11-13) employ different algorithms while sharing a similar baseline-dependent statistical framework for assigning the probability of locus-level and sample-level MSI status. As to the former, for instances, bMSISEA (11) used 100 white blood cells (WBCs) to establish an MSS baseline for each locus, Willis et al. 2019 (5) employed baseline in the process to determine the locus-level threshold through healthy donor samples, and Wang et al. 2020 (13) relied on MSS blood samples as baseline in the locus selection step; sample-level models combine locus-level information and are typically based on a simple threshold for the fraction of unstable loci. The current method also implements a baseline-dependent locus-level confidence framework and unstable locus fraction-based sample-level calling strategy, meanwhile further focusing on subtle aspects that are fundamental to the performance of any cfDNA-based MSI calling algorithm but not explicitly addressed by published methods. Examples are how variation in the empirical duplication level of samples used to construct the baseline could undermine the calling accuracy, and whether the base quality-based deduplication algorithm principally designed for other variants also behaves as anticipated on long mononucleotide repeats, the base quality to the 3’ side of which drops significantly (14,15).

Specifically, we present bMSI-CAST, a novel and ultrasensitive NGS-based, tumour-only cfDNA MSI detection method featuring three key advances: 1) a suite of locus selection rules that can be applied to any given panel to yield mononucleotide loci between 10-13 bp, yields uncompromised base quality and is potentially compatible with all sequencing platforms, with validation for four sequencers from both Illumina and BGISEQ; 2) an MSI-specific duplicate removal strategy that is better suited for mononucleotide locus detection than is Picard MarkDuplicates; and 3) a duplication level-controlled baseline allele frequency distribution to support a Kullback–Leibler divergence (KLD)-based locus-level caller. We sought to provide an accurate and reproducible method for cfDNA MSI detection in clinical cancer decisions, and technical insights regarding key aspects of MSI algorithms.

## MATERIAL AND METHODS

### Study design and sample information

A total of 259 cancer patients and 108 healthy donors were recruited from March 2019 to June 2021 and had their specimens obtained by GeneCast Biotechnology (Supplementary Table 1). Oral and written consent was obtained from the study participants or their representatives prior to enrolment. The studies were approved by the First Affiliated Hospital of Kunming Medical University Ethics Committee (2017-L3). cfDNA and white blood cell (WBC) were obtained from 173 and 198 individuals, respectively. Tumour tissue specimens were available from 149 patients. In details, 148 WBC samples and 40 tumour tissue samples from patients with 8 cancer types (20 positive/20 negative determined by MSI-PCR) were used exclusively for microsatellite selection. To assess the locus-level baseline distribution, we used 70 cfDNA samples. For analytical accuracy assessment, we used 4 previously established MSI-high (MSI-H) cell lines (22Rv1 LACAP RL952 DLD1, COBIOER Inc., China). For clinical validation, we used plasma cfDNA from 103 samples from patients with gastrointestinal cancers. Tumour tissue specimens for all 103 samples were tested by either MSI-PCR or IHC; of these, the tissue, BC and cfDNA samples of 50 positive ones were (4, 11, 29, and 4 in disease stages I-IV, respectively) further sequenced and analysed through an in-house pipeline (see the ‘Bioinformatic pipeline’ section for details) to call point mutations and calculate maximum variant allele frequencies (VAFs). To investigate how coverage varies between sequencing platforms with changing repeat lengths, an independent set of 6 tumour tissue samples was used.

### Sample processing

MSI-PCR was performed using the MSI Analysis System (Promega, USA). Regarding IHC, antibodies used were PMS2, MLH1, MSH6 and MSH2 (ZSGB-BIO, China) and Autostainer Link 48 (Agilent, USA) was used for staining. For the cell line experiment, a fixed amount of 35ng gDNA was sonicated and then titrated with NA12878 (COBIOER Inc., China) to five levels (0.13%, 0.25%, 0.5%, 1.0% and 2.0%) with five technical replicates each. cfDNA was extracted from plasma samples with MagMAX (Thermo Fisher, USA) per the manufacturer’s instructions. Target enrichment was performed with the HyperCap Target Enrichment Kit (Roche, Switzerland) following the manufacturer’s protocol. A fixed mass of products from 8 cycles of pre-capture PCR were fed into 14 cycles of post-capture PCR. The locus selection step was performed based on a 2.09 Mbp, 543 gene pan-cancer panel using tissue specimens sequenced to an average depth of 871. Afterwards, a 100 Kbp gastrointestinal cancer specific panel was designed including selected MSI loci and used to enrich all cell lines gDNA and cfDNA samples used in this study. The resulted libraries were sequenced to an average depth of 25144 (95% CI 5890-46069), corresponding to an average duplication ratio (dup ratio) of 0.84 (95% CI 0.56-0.93). In addition to locus selection, the 2.09 Mbp panel was used for targeted enrichment of tissue samples involved in the estimation of tumour content. All libraries were sequenced in 2×100 paired-end mode on MGI-T7 (BGI-SEQ, China). The 6 tumour tissue samples used for comparison among sequencers were prepared and sequenced by four sequencers: 3 were sequenced by NovaSeq6000 and NextSeq500 sequencers (Illumina Inc., USA) and the other 3 were sequenced by MGI-T7 and MGI-2000 sequencers (BGI-SEQ, China) with identical experimental conditions.

### Bioinformatic pipeline

For both plasma cfDNA and tissue gDNA samples, adaptors were trimmed from raw read pairs using Trimmomatic (v0.36) (16). Clean reads were mapped against the human reference genome (build hg19, UCSC), aligned using BWA bwa-mem (v0.7.12) (17) and sorted using SAMtools (v1.7) (18). A detailed comparison of the majority voting-based deduplication strategy applied by bMSI-CAST and the strategy used by Picard MarkDuplicates (v2.1.0) is described in the “Duplicate removal” section below. MarkDuplicates was performed under the default mode, followed by local indel realignment using GATK version v3.7. For each microsatellite locus, all fragments spanning the locus, defined as those fragments fully covering the microsatellite plus 2 bp in both the 5’ and 3’ directions, were extracted from the realigned BAM file. Following deduplication, the repeat lengths of each locus were assessed and tallied for KLD calculation. Twice the maximum VAF of tumour-informed SNVs called in plasma cfDNA was used to approximate the circulating tumour DNA (ctDNA) content of the clinical samples. The calling methods and filtering criteria are as previously described (19). If no variant was reliably called or the maximum VAF was below the analytically determined LOD (refer to the ‘Results’ section for details), the ctDNA content was considered too low, and the sample was labelled unevaluable.

### Microsatellite locus selection

We adopted a set of filtering steps to screen for high-performance microsatellites: 1) Loci with 10 to 13 mononucleotide A or T repeats were selected. 2) The degree of resemblance of each microsatellite sequence to its immediate upstream or downstream sequences was indicated as the Resemblance Score (RS), which was calculated as

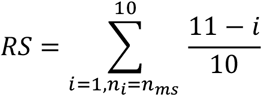

where *n*_*i*_ is the i-th nucleotide counted from either end of the locus outwards, and *n*_*ms*_ represents the repeat unit nucleotide of the target microsatellite locus, counting only those *n*_*i*_ identical to *n*_*ms*_. High RS score indicates poor mapping around the ends of a MSI locus which was discarded if the RS in either direction exceeded a threshold of 2.0. 3) A locus was defined as monomorphic if more than 95% of the group of 118 WBC samples had a central peak allele frequency greater than the locus-specific threshold (longer repeats have lower thresholds). and 4) Loci were sorted by ranked importance via the rank sum test of 20 MSI-H and 20 microsatellite stable (MSS) tissue samples, and loci with p < 0.05 were retained.

### Duplicate removal

Duplicate removal was performed among members of PCR duplications constituting a family with the following criteria. First, only properly paired repeat-spanning fragments were kept. Second, families were defined by the start and end positions. Third, the CIGAR values of all members were inspected to enumerate the reference allele and the insertion and deletion variations. Fourth, the mode was chosen as the final called allele, and the whole family was discarded if there were more than one mode.

The dup ratio of a sample was calculated as a metric to represent the overall family size per locus as follows:

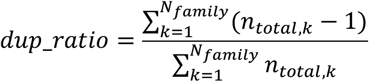

where k denotes the index of a total of *N*_*family*_ families covering the locus, and *n*_*total,k*_ is the member size of family k.

## RESULTS

### Applying the set of microsatellite locus selection rules in the current study

One feature of the current method is a set of locus selection rules aimed to improve the overall accuracy via two mechanisms, namely how easily the loci manifest as *in vivo* MSI-H events, and how significantly the noise level of loci is influenced by *in vitro* PCR processes. In this study, we demonstrated the application of this rules to a given 2.09 Mbp panel.

As to the first mechanism, mononucleotide repeats are prone to somatic variations in cases of MMR deficiency as compared to dinucleotides or longer repeat units, and are therefore preferred and widely chosen. Loci with fewer than 7 repeats are less sensitive to signals of MSI than longer repeats (9,20-22). To further refine the loci, we selected only those loci showing significant KLD scores (P < 0.05, rank sum test) in 20 MSI-H and 20 MSS samples tested by MSI-PCR.

As to the second mechanism, considering differences among nucleotides, A/T repeats are known for being less mutable during PCR than C/G repeats (23). Furthermore, we conducted experiments to further explore possible drawbacks associated with using longer repeats, and we found that mononucleotide repeat length was positively correlated with noise ratio, defined as the sum of the ratios of deleted and inserted alleles to total allele reads (**Fig. 1**). As longer simple repeats are known to be associated with lower base quality at the 3’ end, to evaluate how this could affect MSI calling as a function of locus length and sequencing platform, we further assessed variation in locus coverage over a range of repeat lengths using an independent set of 6 tissue MSS samples on four typically used sequencers from two platforms (MGI-T7 and MGI-2000 from BGI-SEQ and NovaSeq6000 and NextSeq500 from Illumina) (**Supplementary Fig. 1**). Retrievable coverages on NextSeq500 dropped to an unacceptably low level around zero beyond a repeat length of 20 bp; this could be explained by the unique optical system employed by NextSeq500, which stops the sequencing reaction once the base quality drops below a built-in threshold.

**Figure 1.**
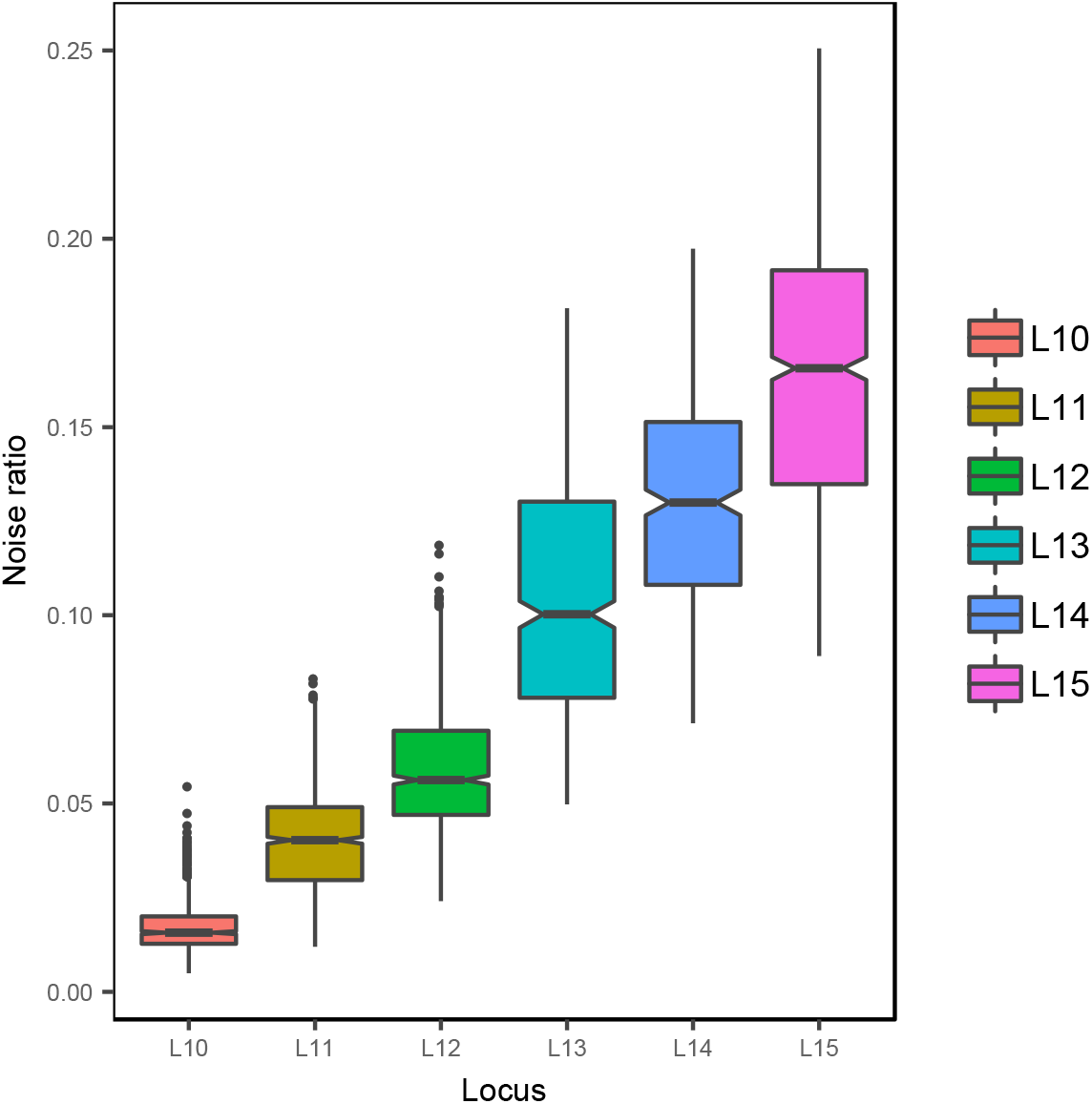
The ratio of the sum of deleted and inserted alleles positively correlates with mononucleotide repeat length. Box plot showing noise ratios (ratio of the sum of deleted and inserted alleles) calculated at six mononucleotide loci (10-15), each with 118 points representing WBC samples from patients with solid tumours. The within-group mean and variance are indicated by the middle bar in boxes and whiskers that correspond to 1.5 interquartile range (IQR).

Considering the aforementioned factors and the raw number of total available loci given the panel size of the current study, it was deemed optimal to use only A/T loci ranging from 10 to 13 bp. A total of 49 mononucleotide loci with high capture efficiency were obtained and used in subsequent stages (**Supplementary Table 2**).

### Comparison of two deduplication strategies

Deduplication is an indispensable denoising step regarding targeted enrichment based NGS analysis. A challenge associated with the accurate deduplication of mononucleotide repeat expansion is that the base quality of sequences flanking the 3’ side of these repeats tends to be lower, while base quality within and upstream of the repeat is not significantly affected, presumably attributable to the phasing/unphasing mechanism associated with the sequencers. Therefore, among reads spanning a locus with an equal number of total bases sequenced, the sum of base qualities associated with longer mononucleotide repeats will theoretically be lower than that of one with shorter repeats. A mixture of truly and falsely deduplicated alleles called from families covering the same microsatellite loci are often tallied in a histogram, constituting a so-called stutter. A deduplication strategy that chooses a read with the greatest sum of sequenced base qualities (GSBQ strategy), such as Picard MarkDuplicates, will be biased toward alleles with shorter repeats and thus result in a left-tilted stutter. In contrast, any algorithm that authentically removes duplicates should result in the reduction of both inserted and deleted allele ratios in the post-deduplication stutter compared with the corresponding pre-deduplication stutter (**Table 1**). To specifically deal with mononucleotide repeats, a simple majority voting duplicate removal strategy was adopted in this study that is predicated on the original repeat length of a locus within a cfDNA fragment remaining the dominant allele throughout library preparation as well as sequencing PCR. To verify the rationality of our approach, the GSBQ and majority voting strategies were compared at six loci (**Fig. 2**). The proportions of total reads represented by the original allele (central ratio), deletion alleles (deleted ratio) and insertion alleles (inserted ratio) were calculated to better characterize the differences. Without deduplication, all three metrics remained unaffected by dup ratio changes, as expected. When the dup ratio increased, the majority voting strategy resulted in a more tightly positioned stutter, evidenced by decreased deleted and inserted ratios and an increased central ratio. This was expected, as an increased duplication level should decrease the noise ratio and thus help better identify the authentic allele. In contrast, Picard MarkDuplicates had a negative impact, i.e., led to worse outcomes than performing no deduplication, as evidenced by a decreased central ratio and an increased deleted ratio. This result was more obvious in longer mononucleotide repeats. The result demonstrated that the majority voting strategy used by bMSI-CAST is more appropriate for the deduplication task on mononucleotide repeats.

**Table 1.** Theoretical dependence of central ratio, deleted ratio, and inserted ratio on increasing the dup ratio using three duplicate removal strategies. Theoretically if duplicates are authentically removed, the central ratio, deleted ratio, and inserted ratio are predicted to be ranked in the order of majority voting < no deduplication < sum of base quality scores; sum of base quality scores < no deduplication < majority voting; and majority voting ∼= sum of base quality scores < no deduplication, respectively.

| Strategy | Deleted ratio | Central ratio | Inserted ratio |
| --- | --- | --- | --- |
| No deduplication | - | - | - |
| Sum of base quality scores (GSBQ) | Increasing | Slightly decreasing | Decreasing |
| Majority voting | Decreasing | Increasing | Decreasing |

**Figure 2.**
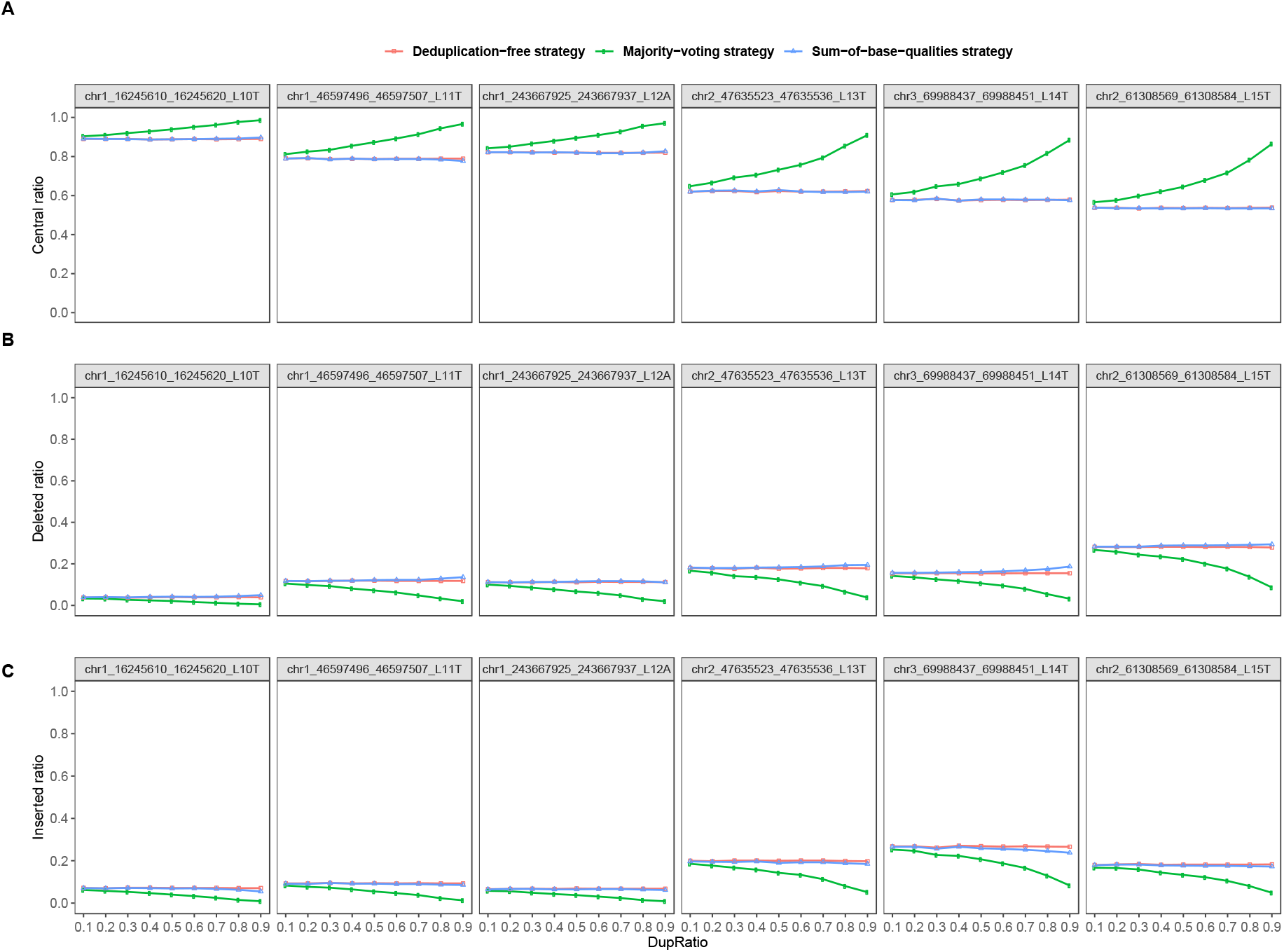
Comparison of three deduplication strategies showed that the majority voting strategy (MV) was more appropriate for deduplication at microsatellite loci with mononucleotide repeats than the greatest sum of base quality (GSBQ) strategy applied by Picard MarkDuplicates. Central ratios (A) deleted ratios (B), and inserted ratios (C) calculated with three deduplication strategies, namely, a sum-of-base-quality strategy by Picard MarkDuplicates (blue), a majority-voting strategy (green) and no deduplication (red). A range of dup ratios were created by the down-sampling of one plasma sample for six representative loci.

### MSI calling algorithm development

Following deduplication, the locus-level MSI status was determined through a statistical test by comparing to a dynamic baseline depending on the duplication level as measured by the dup ratio. The KLD is defined as the relative entropy between two distributions and was used here as the metric to quantify the difference between the allele distributions associated with a locus. Specifically, the KLD value (24) between the allele frequency distribution of a test sample *p*(*x*) and that of the baseline *q*(*x*) was calculated:

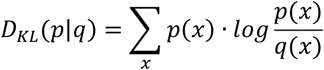

where *p*(*x*) and *q*(*x*) are the frequencies of a given allele *x*.

During baseline construction, *q*(*x*) was constructed with a group of 35 MSS samples as detailed below. Because allele distribution is a function of dup ratios, we built duplication-level-controlled baselines. First, 70 samples were down-sampled to a grid of dup ratios from 0.1 to 0.99 with a step size of 0.01, randomly divided into two equivalent groups of 35 samples, hereby referred to as group A and B, and screened for polymorphisms. At each dup ratio level, the allele frequencies of each observed allele were averaged across group A, resulting in a vector of means *q*(*x*). The KLD score of each sample in group B was calculated with respect to *q*(*x*), which represented the background level distribution and was fitted by a gamma distribution. This resulted in a set of null distributions for individual loci. As the dup ratio increased from 0.5 to 0.9, the Central ratio and noise ratio of the baseline increased and decreased, respectively (**Supplementary Fig. 2**).

Given a test sample, the dup ratio was first calculated for each locus. Second, the corresponding dup-ratio-matched baseline allele frequency distributions were used to calculate KLD scores and any locus with a KLD score falling outside the upper limit of the 99.5% confidence interval was considered unstable at that specific locus. Third, for sample-level MSI status calling, the fraction of unstable loci was compared to the threshold, and the sample was called MSI-H or MSS accordingly, where a minimum of 15 loci were required. In summary, the locus selection rules, the majority voting strategy, and the KLD method with dup ratio-controlled baselines constitute the bMSI-CAST.

### Analytical and clinical validation of bMSI-CAST

To analytically assess the performance of bMSI-CAST, we first set a lower threshold for MSI positivity. We performed MSI scoring for 70 cfDNA samples from healthy donors; the maximum obtained value of 0.13 was used as the threshold in subsequent analyses (**Fig. 3A**). Analytical sensitivity was then evaluated using four established cell lines, diluted to five levels from 0.13% to 2% with 20 data points for each titration level (19 for 0.13% due to technical failure in 1 experiment). The sensitivity was 79% (15/19) at a minimum of 0.13% and 100% for higher concentrations. The 95% LOD was determined to be 0.15% through Probit regression (**Fig. 3B**).

**Figure 3.**
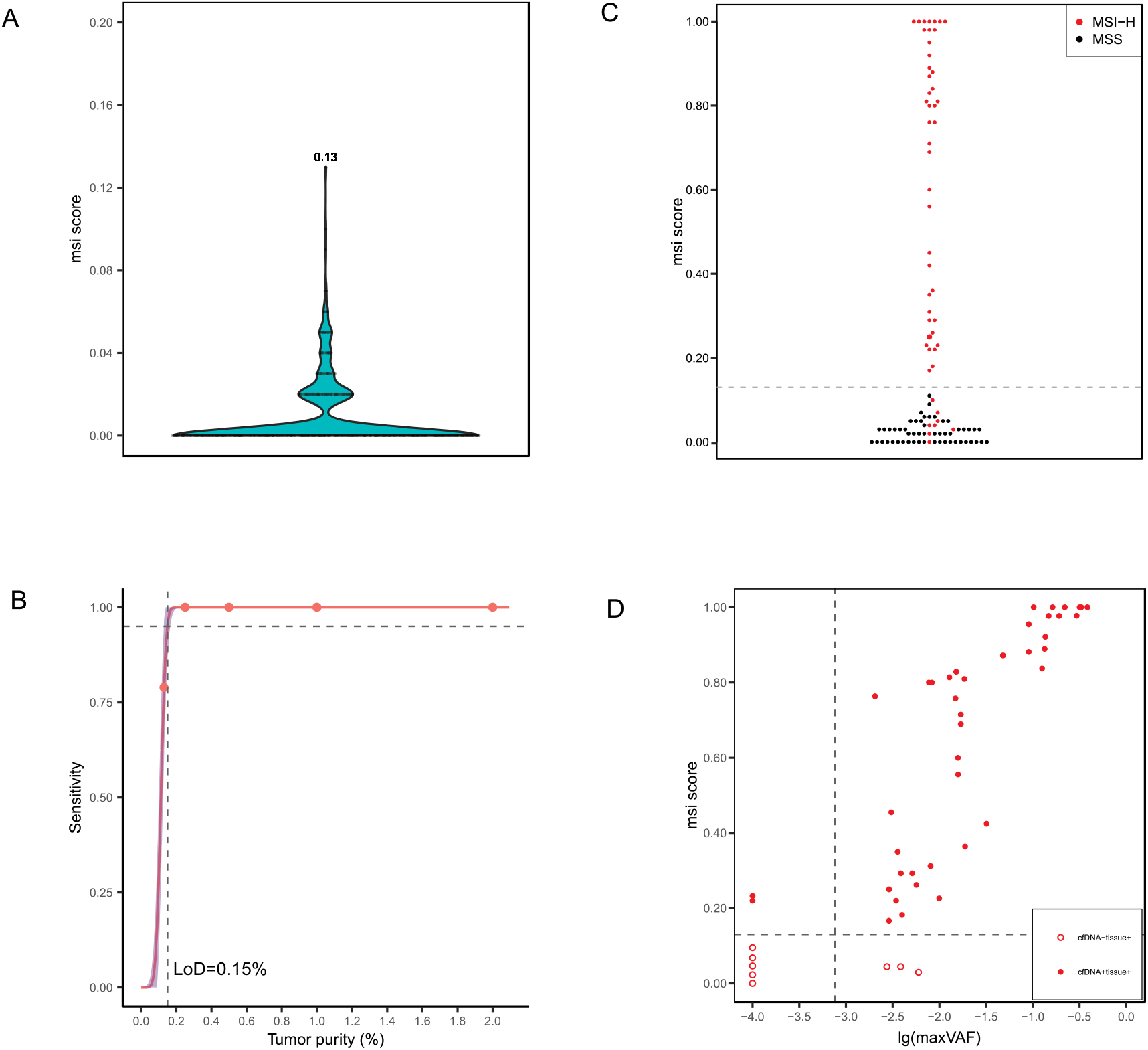
Determination of the MSI score threshold. (A) Violin plot showing the MSI score distribution for 70 cfDNA samples from healthy donors. 0.13 was the maximum MSI score. (B) Determination of the analytical sensitivity of bMSI-CAST. Probit regression performed on sensitivities calculated on four diluted cell line samples (22RV1, DLD1, LNCAP, and RL952) at five ctDNA content levels (0.13%, 0.25%, 0.5%, 1.0% and 2.0%, each with four to five replicates); the 95% confidence interval is shown by purple shading. The dashed line indicates the 95% LOD that corresponds to 0.15% tumour purity. (C) Clinical validation of bMSI-CAST. Beeswarm plot showing the MSI score distribution for 103 clinical cfDNA samples (50 positive and 53 negative). The dash line indicates the analytically determined MSI score threshold of 0.13. (D) Concordance of bMSI-CAST results with tissue MSI-H results for 102 clinical cfDNA samples, with one sample excluded due to a lack of ctDNA content information. Solid and open circles indicate MSI-H and MSS called by bMSI-CAST, respectively. The vertical dash line at 0.075% indicates the threshold for evaluable samples., and the horizontal dash line indicates at 0.13 indicates the threshold for MSI score.

The clinical accuracy of bMSI-CAST was evaluated by comparing the classification results of 103 cfDNA samples from patients with gastrointestinal to those of tissue MSI-PCR. Eight of the samples (8/103, 10%) had no somatic variant with VAF >= 0.2% and were considered unevaluable for MSI status (5). Among the 95 evaluable samples (95/103), bMSI-CAST exhibited a positive predictive agreement (PPA) of 92.9% (39/42, 95% CI 79.4∼98.1%) and negative predictive agreement (NPA) of 100% (53/53, 95% CI 92∼100%) with tissue MSI-PCR, for an overall accuracy of 96.8% (92/95, 95% CI 90.3∼99.2%) and a positive predictive value of 100% (39/39, 95% CI 88.9∼100%) for patients across all clinical stages (**Fig. 3C and Fig. 3D**). Moreover, 3 of the 8 unevaluable MSI samples could be assessed by bMSI-CAST (data not shown). Together, these results demonstrate the excellent performance of bMSI-CAST and its promise for diagnosing MSI status from cfDNA samples.

## DISCUSSION

We have characterized the three main features of bMSI-CAST and validated the overall performance. The set of locus selection rules had multiple factors taken into consideration and was aimed to achieve both high accuracy and compatibility across all sequencing platforms. Furthermore, the majority voting deduplication strategy was a better choice for deduplication regarding mononucleotide repeats than the GSBQ strategy used by Picard MarkDuplicates. The use of duplication-level specific baselines in combination with a baseline-dependent metric, KLD score, was also demonstrated.

The proposed locus selection principles address a comprehensive list of factors, some of which were not previously considered or explicitly formulated. First, in the absence of a patient-matched normal sample, polymorphic loci ought to be screened for specificity (6,25,26), optimally using samples from the same population (27); in this study we used a set of WBC samples all from the Chinese population to select monomorphic loci. Note that if the patient-matched normal is available, it can be used in addition. Second, compatibility of MSI loci across sequencing platforms has not been assessed by existing methods. Here we demonstrated the association between the retrievable coverage and the repeat length on 4 sequencers from 2 platforms. Given the panel size of 2.09 Mbp in this study, 49 loci with size between 10 to 13 were selected and deemed enough. In general, we recommend using loci shorter than 15 bp which is presumably compatible with all NGS sequencing platforms as discussed above. To further improve the overall performance, the thresholds associated with selection rules can be customized to fit user specific scenarios and requirements. For example, choosing a RS threshold greater than 2.0 will result in a greater number of but less specific loci and vice versa. Furthermore, a larger panel or search scope in the genome can provide leeway for applying more stringent threshold, or otherwise there will be too few loci to achieve required sensitivity. Last but not least, the number of loci influences the overall accuracy intricately through a trade-off that including more loci can in theory continuously increase sensitivity, whereas excluding nonspecific loci can help improve specificity in any algorithm that is contingent on the fraction of unstable loci. Using more than 40 samples to select significant loci will also help further improve the performance. In summary, we have explicitly formulated a set of locus selection rules aimed to both be practical and include as many relevant factors as possible.

Published NGS-based MSI detection methods have either explicitly stated that they used Picard to remove duplicates (4,28) or not mentioned whether or how duplicate removal was performed (8,12,29). Picard MarkDuplicates chooses the read with the greatest sum of sequenced base qualities and has been widely used in non-MSI applications. To the best of our knowledge, whether this strategy is suitable for reads covering long mononucleotide repeat sequences has not been assessed. Deduplication is fundamentally a process to discriminate true signal from noise, and its accuracy is a function of the dup ratio, repeat locus characteristics, and deduplication strategy used. We have showed the superiority of our majority voting strategy for deduplication over a GSBQ strategy for 10 to 13 bp mononucleotide loci. Independent of bMSI-CAST, the majority voting strategy can be universally applied to duplication of mononucleotide repeats.

Implementing a duplication level-dependent baseline demands an approximate and accurate enough way to abstract the real duplication level. Clinical cfDNA samples inevitably vary in terms of input masses, which in turn affects the number of PCR cycles to run and the library complexity; the analysis of these samples is further complicated by varying sequencing depths. The dup ratio serves as a metric to approximate the combined effect of these variables, and matching test samples to a baseline allele frequency distribution according to dup ratios essentially calibrates the family size distribution of data input to the downstream calling algorithm and contributes to the overall accuracy of bMSI-CAST and arguably any baseline-dependent MSI algorithms. We note that the duplication level-controlled baseline approach described here can potentially be applied to any baseline-dependent MSI calling algorithm that uses a metric other than KLD score, and further improve its performance, and the baseline distribution should be built for a fixed and stable experimental pipeline and inspected for suspicious outliers or abnormally high variances before being applied to test samples. It must be considered that the dup ratio is only an approximation and therefore is not able to represent differences in every single variable. It has been reported that the ratios of deleted or inserted alleles after a number of cycles depend on varying repeat unit length, repeat unit nucleotides, number of repeats and polymerase types (30). It is theoretically possible that two post-PCR libraries with the same dup ratios may correspond to different numbers of PCR cycles. Therefore, to use bMSI-CAST or any other MSI detection tool, one should first evaluate the influences of PCR cycle variations. Throughout the current study, 22 PCR cycles (8 cycles of pre-capture PCR plus 14 cycles of post-capture PCR) were used, far below the 47 cycles threshold after which 10-13 bp long A/T mononucleotide repeats become too noisy to be informative; therefore, we deemed it safe to not correct for PCR influences in the present pipeline. This again stresses the benefit of choosing shorter repeats. Independent of bMSI-CAST, the use of a dup ratio-matched baseline could presumably help improve calling accuracy in any custom pipelines involving comparison to a baseline.

We discuss one main difference in the calling algorithm between bMSI-CAST and other methods here. In theory, MSI events are detectable based on differences in the number of alleles, counts of alleles and order or pattern of the counts of alleles, and then assigning confidence to such differences under assumptions of the signal and noise distribution. Wang et al. 2020 (13) retained repeat lengths less than or equal to a threshold repeat length that provided the greatest difference between a group of MSS samples and a group of MSI samples. Similarly, bMSISEA chose MSI pattern with a range of deleted repeat lengths using long repeat lengths (16-27). Realizing that all deleted and inserted alleles inevitably present both noise and true signals, we utilized the KLD score to consider the whole distribution of repeat lengths and take full advantage of all potential signals, along with using a dup ratio-controlled baseline allele frequency distribution to curtail noise. Because KLD scores can theoretically capture the inherent order information of alleles in a stutter in addition to the abundance of alleles, bMSI-CAST provides more comprehensive characterization of a stutter and higher resolution. These theories were borne out by the strong performance of bMSI-CAST in clinical and analytic validation experiments. Worth noting is that a stringent comparison with published methods was hard due to unavailability of tools or failure to construct baselines. As an indirect comparison of reported performances, Willis et al. reported a 95% limit of detection (LOD) of 0.1% with 30 ng input, and clinical sensitivity and specificity of 87% and 99.5%, respectively, with 90 loci and a lower threshold for maximum VAF of 0.2%. bMSISEA had an LOD of approximately 0.35%, and clinical sensitivity and specificity of 94.1% and 100%, respectively, with a lower threshold for maximum VAF of 0.2% and 8 mononucleotide loci with repeat lengths ranging from 16 to 27 when the sequencing depth was approximately 15000X (11). Wang et al. 2020 reported an LOD of 0.5% for 30 ng input and sensitivity of 82.5% and specificity of 96.2%, with 100 loci and a lower threshold for maximum VAF of 0.5% (13). bMSI-CAST showed an LOD of 0.15%, and clinical sensitivity and specificity to be 92.9% and 100%, respectively, with 49 loci and a lower threshold for maximum VAF of 0.2%, parallel to or outperforming the peer methods.

As to possible future improvement, instead of retrospectively selecting loci given an existing panel, the set of locus selection rules make it possible for a search in the whole exome or even the whole genome to prospectively select loci for better performance under more stringent criteria. Also, bMSI-CAST should be straightforward to use in combination with UMI, which we expect will further increase the overall performance. In conclusion, we have developed and validated a novel plasma cfDNA-based MSI detection method to help better aid non-invasive diagnosis and prognosis of cancer. We anticipate that existing methods could have benefited from incorporating the three features into their specific algorithms. Next, we plan to validate bMSI-CAST in a larger cohort and investigate the responsivity to immunotherapy in prospective interventional studies.

## DATA AVAILABILITY

bMSI-CAST can be applied as a complete tool, or the integral parts other than the calling algorithms can be combined with a chosen calling algorithm in general, all of which are freely accessible at https://github.com/GeneCast/bMSI-CAST.

The raw sequence data reported in this paper have been deposited in the Genome Sequence Archive (31) at the National Genomics Data Centre (32), Beijing Institute of Genomics (China National Centre for Bioinformation), Chinese Academy of Sciences, under accession number subHRA000493, which is publicly accessible at http://bigd.big.ac.cn/gsa-human.

## SUPPLEMENTARY DATA

Supplementary Data are available at NAR online.

## ACKNOWLEDGEMENT

This work was supported in part by the National Natural Science Foundation of China No. 31660312. The authors thank the patients and staff at the First Affiliated Hospital of Kunming Medical University.

## TABLE AND FIGURES LEGENDS

**Supplementary Figure 1.**
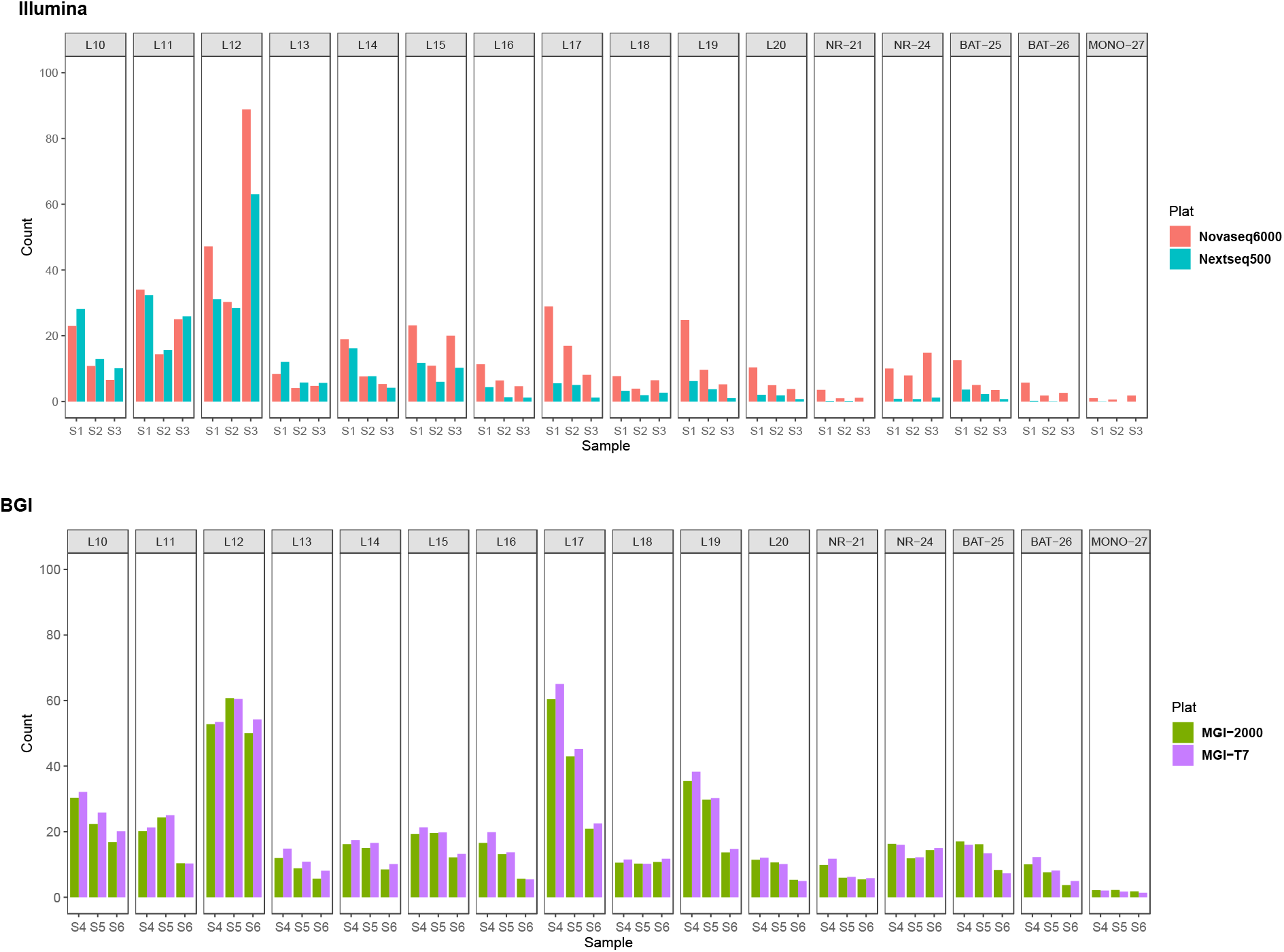
Coverage values over 16 mononucleotide loci with repeat lengths ranging from 10-27 were compared between two pairs of sequencers: the two Illumina platform sequencers NovaSeq6000 vs. NextSeq500 and the two BGI platform sequencers MGI-2000 vs. MGI-T7.

**Supplementary Figure 2.**
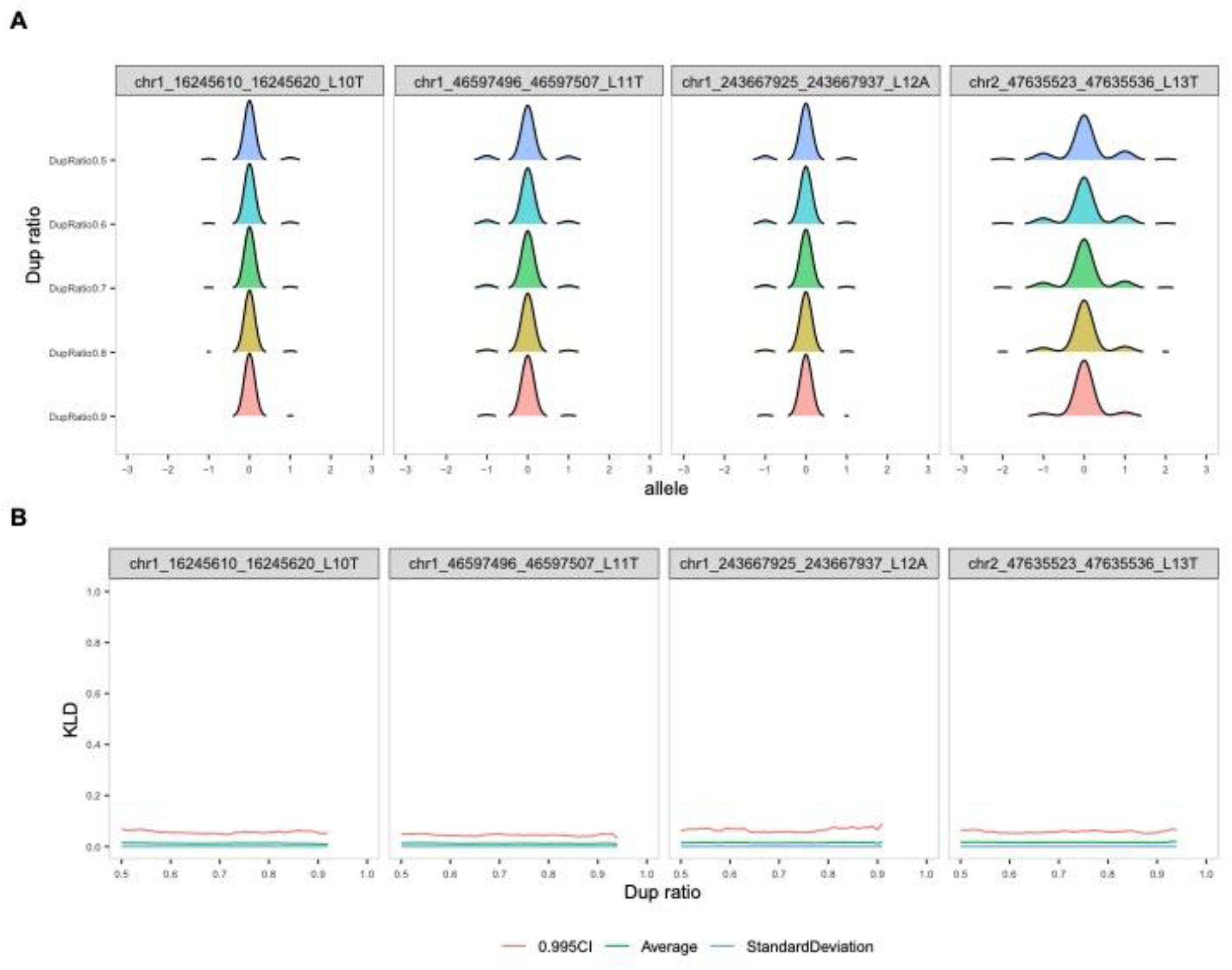
Characterization of the duplication level-matched baseline approach. (A) Ridgeline plot showing how the central and noise ratios change as the dup ratio (0.5 to 0.9) and the repeat length (10 to 13) increases. Zeros on the x axis indicate the central allele, and the positive and negative offsets represent inserted and deleted alleles, respectively. (B) For the same dup ratio range and discrete repeat lengths as in (A), the KLD scores were calculated for a grid of dup ratios with step size of 0.01 and plotted as a smooth line (green), where values corresponding to the 99.5% CI and the standard deviation were also plotted (red and blue).

Supplementary Table 1 Detailed sample information.

Supplementary Table 2 Detailed locus information.

